# Ares-GT: design of guide RNAs targeting multiple genes for CRISPR-Cas experiments

**DOI:** 10.1101/2020.01.08.898742

**Authors:** Eugenio G. Minguet

**Affiliations:** Independent Researcher, Valencia, Spain

## Abstract

**Motivation:** There is a lack of tools to design guide RNA for CRISPR genome editing of gene families and usually good candidate sgRNAs are tagged with low scores precisely because they match several locations in the genome, thus time-consuming manual evaluation of targets is required. Moreover, online tools are limited to a restricted list of reference genome and lack the flexibility to incorporate unpublished genomes or contemplate genomes of populations with allelic variants.

**Results:** To address these issues, I have developed the ARES-GT, a local command line tool in Python software. ARES-GT allows the selection of candidate sgRNAs that match multiple input query sequences, in addition of candidate sgRNAs that specifically match each query sequence. It also contemplates the use of unmapped contigs apart from complete genomes thus allowing the use of any genome provided by user and being able to handle intraspecies allelic variability and individual polymorphisms.

**Availability:** ARES-GT is available at GitHub (https://github.com/eugomin/ARES-GT.git).

---

The design of optimal single guide RNAs (sgRNAs) is a critical step in CRISPR/Cas genome editing, and it must ensure specificity and minimize the possibility of offtarget mutations. Although good online tools are available for identification of CRISPR DNA targets, which have popularized genome editing, their use is limited to a restricted list of genomes (Bae et al., 2014;Heigwer et al., 2014;Lei et al., 2014;Haeussler et al., 2016;Liu et al., 2017;Labun et al., 2019), sometimes corresponding to less than ten species (Pliatsika and Rigoutsos, 2015;Doench et al., 2016). Even Breaking-Cas (Oliveros et al., 2016), a free online tool which currently offers more than 1600 genomes, lacks the flexibility to easily incorporate unpublished genomes or contemplate genomes of populations with allelic variants -an issue partially addressed by AlleleAnalyzer for the human genome (Keough et al., 2019). An additional problem posed by the design of sgRNAs targeting gene families is that good candidate sgRNAs can be tagged with low scores precisely because they match several locations in the genome, thus time-consuming manual evaluation of targets is required. To address these issues, I have developed the ARES-GT, a local command line tool in Python programming language (https://www.python.org/).

ARES-GT can identify targets of the two most widely used CRISPR enzymes (Cas9 and Cas12a/Cpf1) and evaluates possible offtargets in a user-provided reference genome, including non assembled contigs and unpublished genomes from any species. A list is generated with the best candidates (those with no offtargets based on parameters selected by user) and, if multiple query genes from the same family are targeted, the list includes sgRNAs that match more than one of them. Detailed information for each possible target is also provided, including an alignment with the possible offtargets. ARES-GT have been already used succesfully in *Arabidopsis*, tomato and rice while under development (Aliaga-Franco et al., 2019;Bernabé-Orts et al., 2019).

It has been reported that the specificity of both Cas9 and Cas12a is particularly sensitive to mismatches in the PAM proximal sequence (on an 11- and 8-nucleotide stretch for Cas9 and Cas12a, respectively), named “seed” (Cong et al., 2013;Hsu et al., 2013;Zetsche et al., 2015;Swarts et al., 2017). Mismatches in the seed sequence has a critical impact into cleavage efficiency on DNA target, and it is unlikely that seed sequences with 2 or more mismatches cause real offtargets *in vivo. S*equence composition and the number and distribution of mismatches also affects cleavage efficiency (Hsu et al., 2013). Therefore the ARES-GT algorithm discards possible offtargets using as criterium the presence of 2 or more mismatches in the seed sequence, while the user defines the second threshold criterium: the number of total mismatches when there are none or one mismatches in the seed sequence. In addition, the user must also indicate whether a “NAG” PAM, which Cas9 can recognise though with lower efficiency (Hsu et al., 2013), must be taken into account when evaluating possible offtargets.

## Design of guide RNA matching multiple CBF genes

As a proof of concept, I have choosen the C-repeat/DRE-Binding Factor (CBF) gene family of plant transcription factors to test the various novelties implemented in ARES-GT. Among the four members identified in *Arabidopsis thaliana*, three of them –*AtCBF1, AtCBF2* and *AtCBF3*–, have been implicated in the response to cold temperatures, while *AtCBF4* has been implicated in the response to drought (Haake et al., 2002;Yamaguchi-Shinozaki and Shinozaki, 2006). The first three members of this family are closely located in less than 8 Kb in chromosome 4 (Figure 1A), making extremely difficult to obtain a triple mutant by classical crossing strategy. This has been recently achieved by CRISPR/Cas9-induced mutagenesis (Cho et al., 2017) using two sgRNAs that the authors selected by manual evaluation of sequence alignments, manual selection of candidates, and specificity verification with CRISPR-P (Lei et al., 2014). I used the *A. thaliana* genomic coding sequences (TAIR v10) of the four *CBF* genes as a multiple query in ARES-GT, to search for candidate sgRNAs using both Cas9 and Cas12a. A total of 96 and 34 unique specific targets matching only one location in the genome and with no predicted offtargets were found for each the four genes, using Cas9 and Cas12a, respectively. More interestingly, the program also listed 13 candidates for Cas9 and 10 candidates for Cas12a that match multiple *CBF* genes (Tables 1 and 2). In total, 10 Cas9 and 5 Cas12a candidates were identified that match more than one *CBF* gene and did not present any offtarget outside *CBF* genes (Figure 1B, 1C). Among them were included the two sequences previously reported (Cho et al., 2017), corresponding to Cas9CBF1_015 and Cas9CBF2_124 in this work.

**Table 1.**
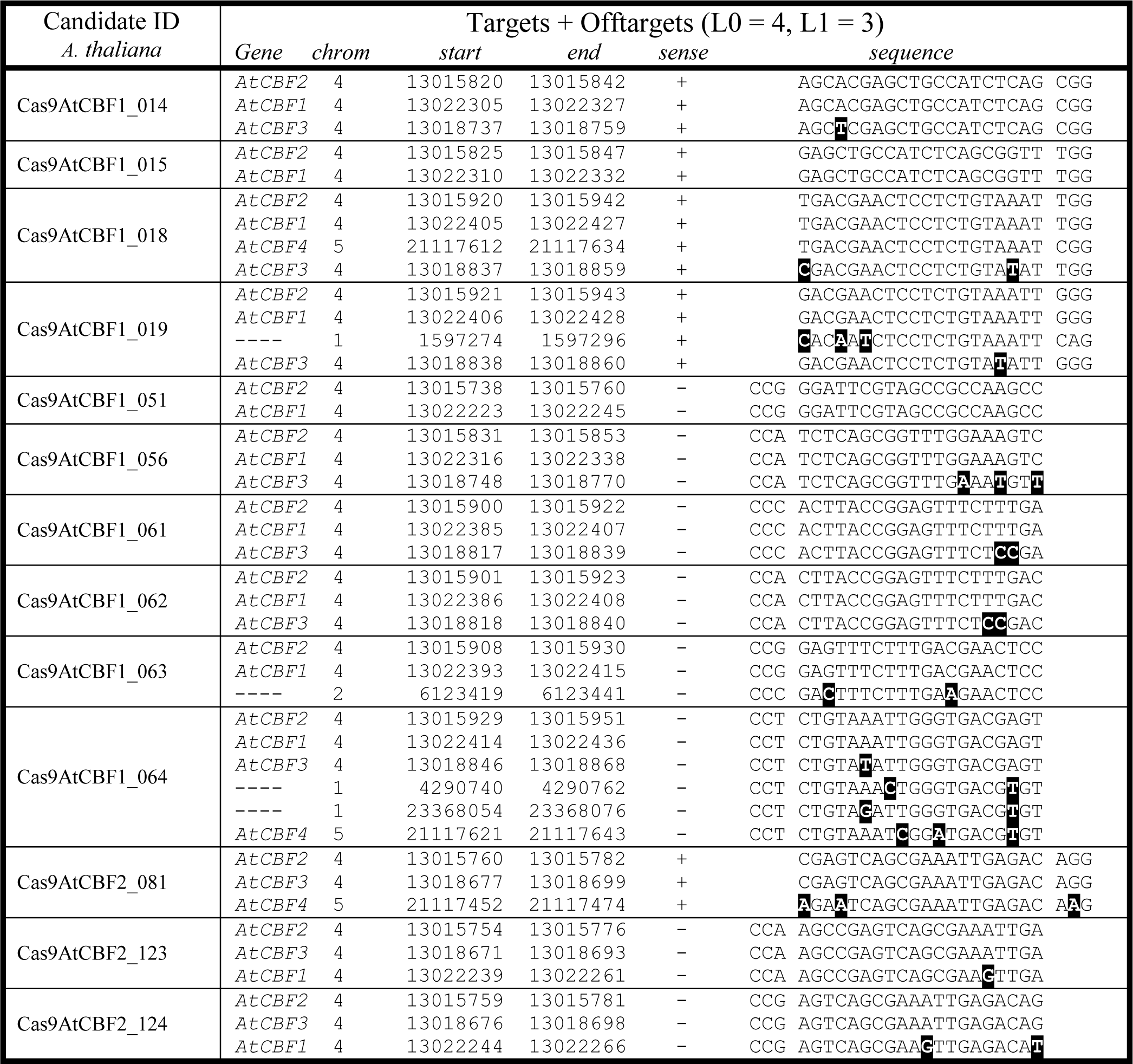
Multiple targets Cas9 candidates for *AtCBF* genes. All possible genome targets and offtargets (with ARES-GT thresholds: L0 = 4 and L1 = 3) of each candidate are listed with indication of genome coordinates (TAIR v10) and whether it corresponds to a *CBF* gene. In alignments, black boxes mark mismatches and a space separates PAM (NGG or NAG) from sequence. Differences in the “N” position in the PAM are not marked.

**Table 2.**
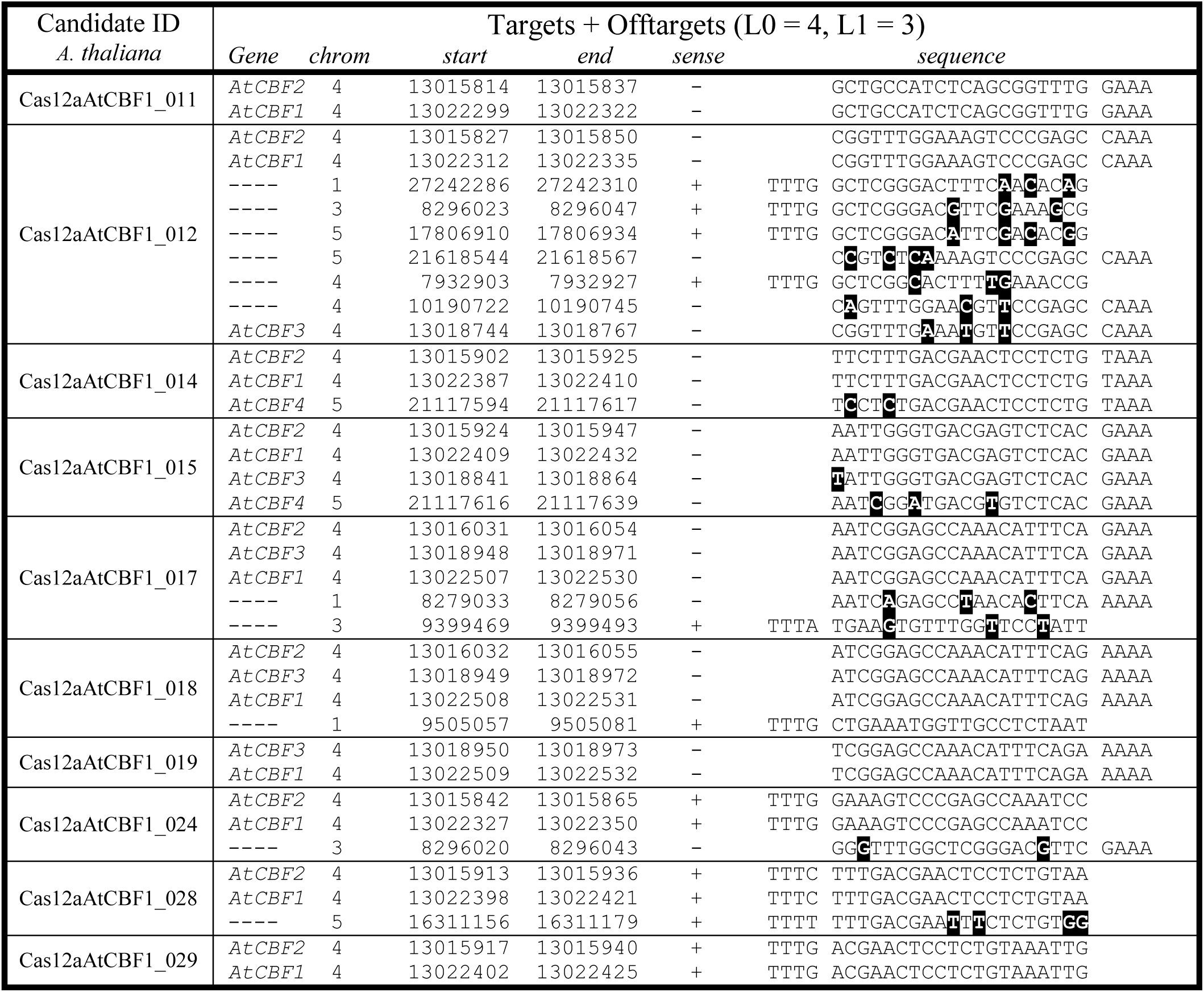
Multiple targets Cas12a candidates for *AtCBF* genes. All possible genome targets and offtargets (with ARES-GT thresholds: L0 = 4 and L1 = 3) of each candidate are listed with indication of genome coordinates (TAIR v10) and whether it corresponds to a *CBF* gene. In alignments, black boxes mark mismatches and a space separates PAM (TTTN) from sequence. Differences in the “N” position in the PAM are not marked.

**Figure 1:**
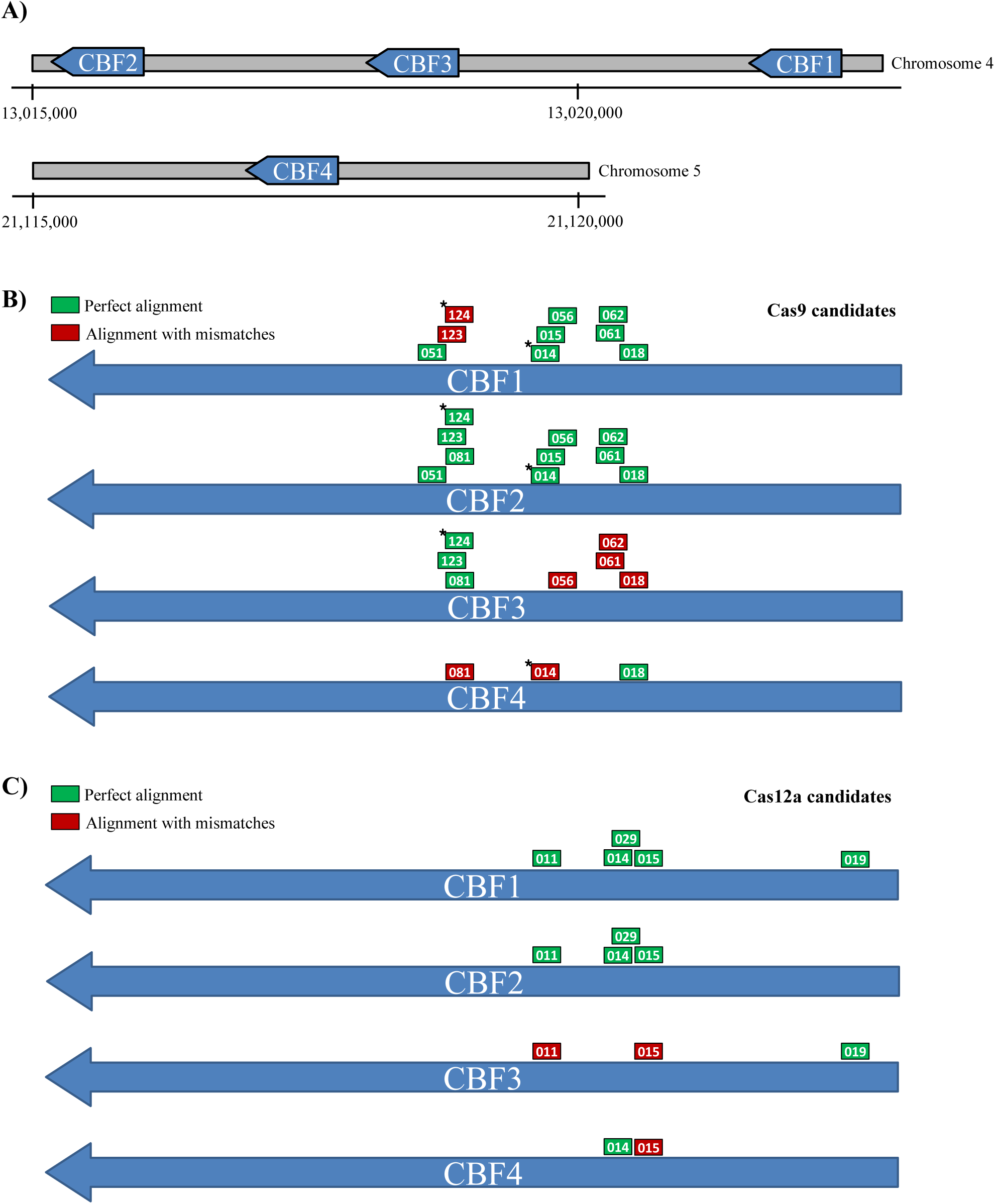
A) Genomic distribution of CBF genes in *Arabiopsis thaliana* chromosomes 4 and 5. Location of Cas9 (B) and Cas12a (C) candidates with multiple CBF gene targets. (*) Asterik marks candidates corresponding with previously reported sgRNAs (Cho et al., 2017).

To test that AREST-GT can work with any user-provided genome, including unmapped contigs, I selected the first version of the genome of *Cardamine hirsuta* (Gan et al., 2016). The available genome sequence spans over its 8 chromosomes, but also contains 622 unmapped contigs in addition to chloroplast and mithocondria genomes. The sequence information was downloaded (http://chi.mpipz.mpg.de/index.html) and used locally with ARES-GT for searching CRISPR targets in the four *C. hirsuta CBF* homologous genes (Supplementary Figure 1). In addition to unique specific targets (86 for Cas9 and 28 for Cas12a), 10 candidate sgRNAs for Cas9 and 3 for Cas12a were identified that perfectly match *ChCBF1* and *ChCBF2* (Table 3). Taking into account possible offtargets, only 5 and 3 sequences for Cas9 and Cas12a, respectively, are relyable candidate sgRNAs targetting only *ChCBF* family genes. For instance, Cas9ChCBF1_044 perfectly matches *ChCBF1* and *ChCBF2*, and it also matches *ChCBF3* with one mismatch.

**Table 3.**
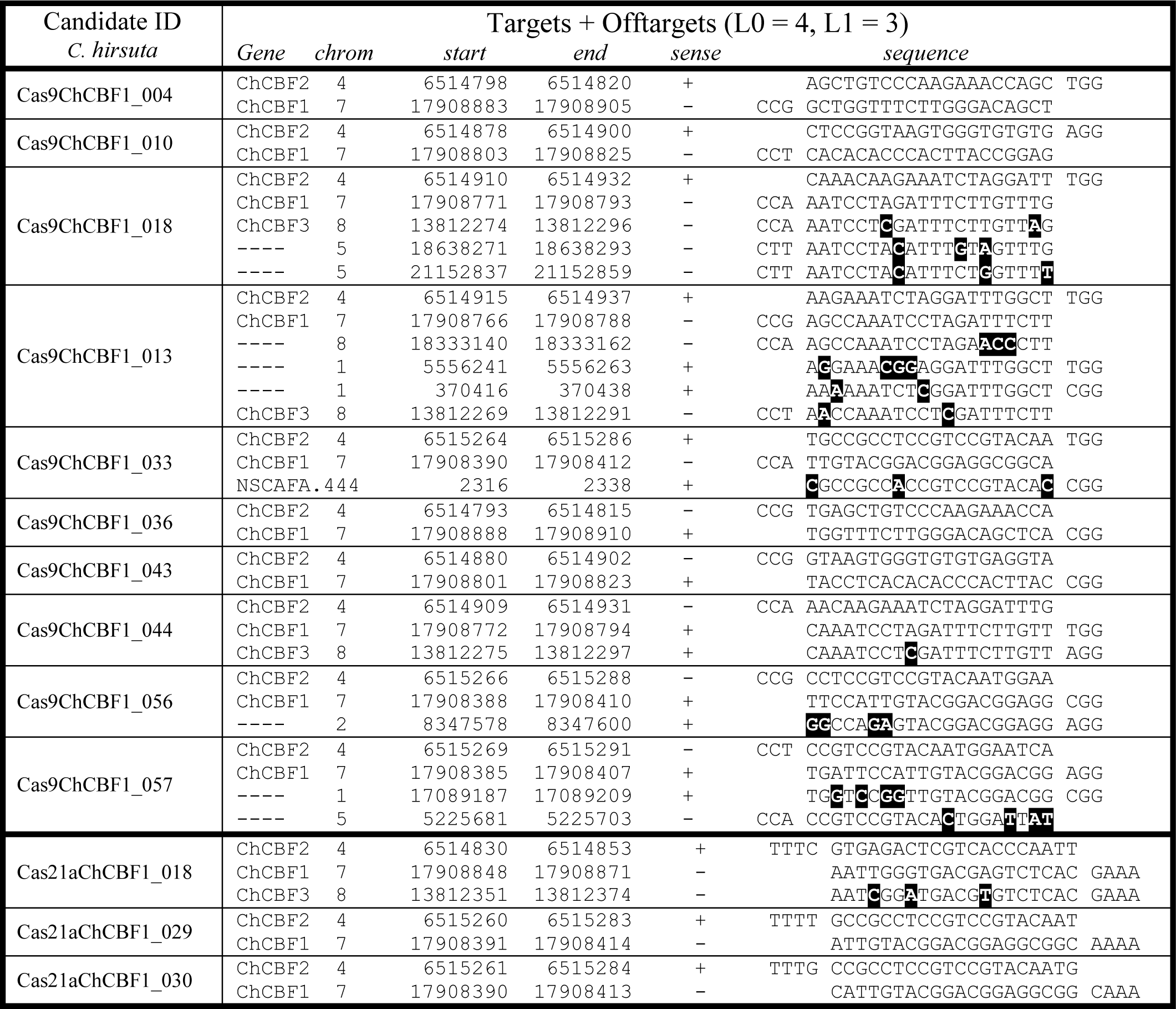
Multiple targets Cas9 and Cas12a candidates for *ChCBF* genes. All possible genome targets and offtargets (with ARES-GT thresholds: L0 = 4 and L1 = 3) of each candidate are listed with indication of genome coordinates (*Cardamine hirsuta* v1.0) and whether it corresponds to a *CBF* gene. In alignments, black boxes mark mismatches and a space separates PAM (NGG/NAG or TTTN) from sequence. Differences in the “N” position in the PAM are not marked.

Finally, to contemplate intraspecific allelic variability in the design of sgRNAs for genome editing, I used ARES-GT in combination with the genome sequences available through the Arabidopsis 1001 genomes project (https://1001genomes.org/). Contrary to available online tools, which only work with the standard *A. thaliana* Col-0 accession, ARES-GT can be used to design ecotype-specific editing tools taking advantage of polymorphic sequences in the different accessions. Good quality genome assemblies of seven *A. thaliana* accessions (*An-1, C24, Cvi, Eri, Kyo, Ler* and *Sha*) (Jiao and Schneeberger, 2019) were downloaded, and ARES-GT was used to design sgRNAs targetting CBF genes in each accession. As reflected in Table 4, the SNPs in *CBF* genes between the different accessions are responsible of the identification of different number of candidate sgRNAs that match several genes of the family, from 18 Cas9 candidates with *CBF* genes from *Kyo* genome to 11 Cas9 candidates with *CBF* genes from *Cvi* genome. The selection of CRISPR candidates with specific unique target (without offtargets) also varied between accessions (Table 4). I used each accession CBF genes as query for ARES-GT but using either the standar *Col-0* reference or the corresponding accession genome. Candidates only listed when *Col-0* is used as reference (*Col-0* exclusive) are false positives, as they have offtargets in the corresponding accession genome. The accession’s exclusive candidates would be false negatives, as they are discarded if *Col-0* is used but do not have offtargets in the corresponding accession genome (Table 4). Differences in the identification of offtargets also affects the selection of efficient candidates matching several CBF genes. For instance, candidate C24_CBF1_019 perfectly match C24_CBF1, C24_CBF2 and C24_CBF3 but has a possible offtarget (4 mismatches in distal sequence) in the chromosome 3 of *C24* genome, which is above offtarget thresholds in *Col-0* genome because of an extra mismatch in the proximal sequence (Table 5). In the other sense, Eri_Cas12aCBF1_017 is a candidate that perfectly match Eri_CBF1, Eri_CBF2 and Eri_CBF3 without offtargets in Eri genome, however it would be discarded because two offtargets are detected if *Col-0* genome is used (Table 5).

**Table 4.**
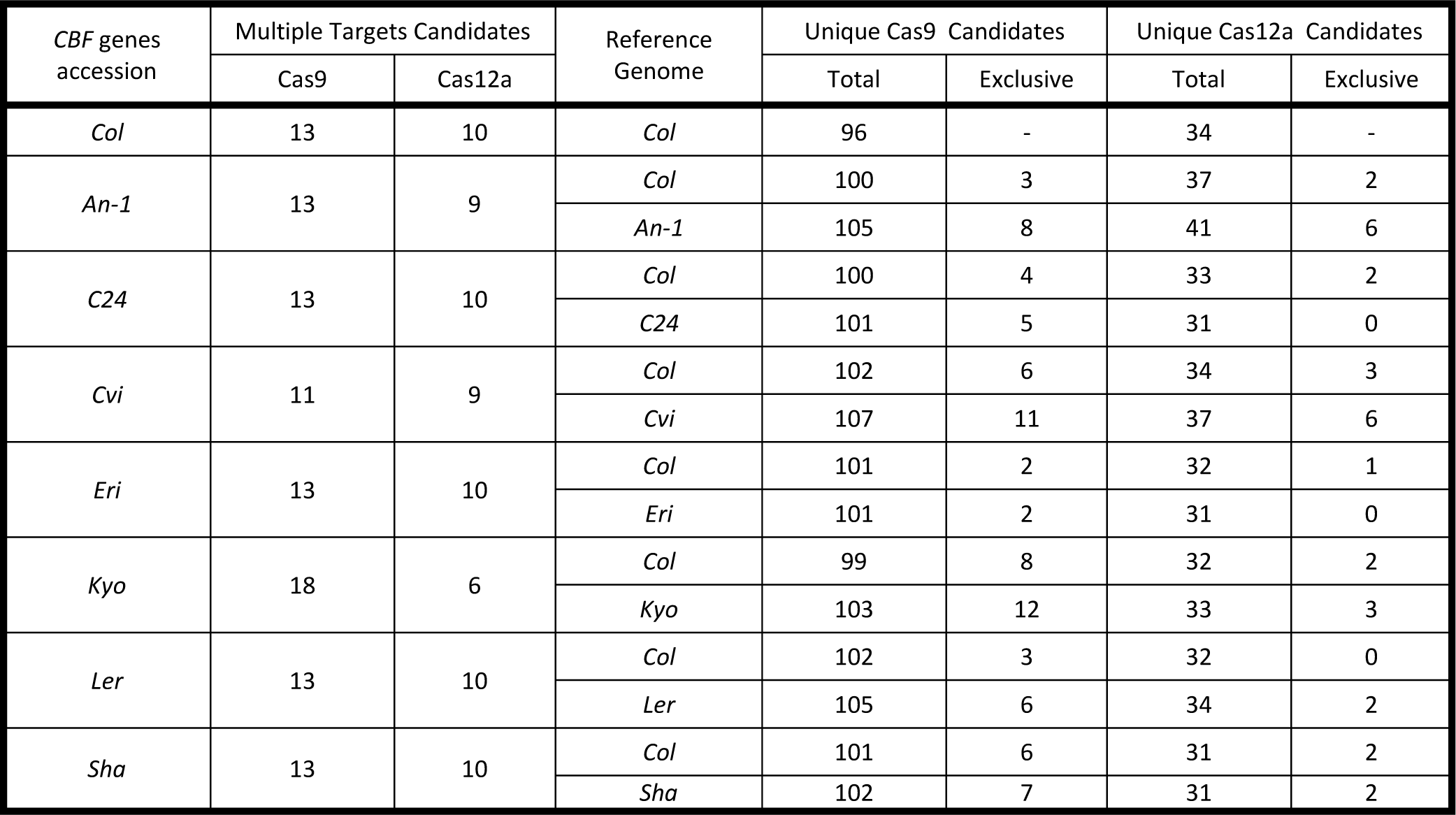
Intraspecies variability effect in the number of Cas9 and Cas12a candidates targetting multiple or unique *AtCBF* genes. Sequence variability in the *CBF* genes from different *Arabidopsis thaliana* accessions change the number of candidates that can match multiple targets due to SNPs in the 20 nucleotides of the guide but also SNPs affecting PAM sequence. The use of the standard *Col-0* genome reference (TAIR v10) or the corresponding accession genome affects the identification of offtargets thus the correct identification of specific (unique) candidates matching only one *CBF* gene. The column “exclusive” indicates the number of specific candidates that are only listed when the corresponding reference genome is used.

| *CBF* genes accession | Multiple Targets Candidates |  | Reference Genome | Unique Cas9 Candidates |  | Unique Cas12a Candidates |  |
| --- | --- | --- | --- | --- | --- | --- | --- |
|  | Cas9 | Cas12a |  | Total | Exclusive | Total | Exclusive |
| *Col* | 13 | 10 | *Col* | 96 | - | 34 | - |
| *An-1* | 13 | 9 | *Col* | 100 | 3 | 37 | 2 |
|  |  |  | *An-1* | 105 | 8 | 41 | 6 |
| *C24* | 13 | 10 | *Col* | 100 | 4 | 33 | 2 |
|  |  |  | *C24* | 101 | 5 | 31 | 0 |
| *Cvi* | 11 | 9 | *Col* | 102 | 6 | 34 | 3 |
|  |  |  | *Cvi* | 107 | 11 | 37 | 6 |
| *Eri* | 13 | 10 | *Col* | 101 | 2 | 32 | 1 |
|  |  |  | *Eri* | 101 | 2 | 31 | 0 |
| *Kyo* | 18 | 6 | *Col* | 99 | 8 | 32 | 2 |
|  |  |  | *Kyo* | 103 | 12 | 33 | 3 |
| *Ler* | 13 | 10 | *Col* | 102 | 3 | 32 | 0 |
|  |  |  | *Ler* | 105 | 6 | 34 | 2 |
| *Sha* | 13 | 10 | *Col* | 101 | 6 | 31 | 2 |
|  |  |  | *Sha* | 102 | 7 | 31 | 2 |

**Table 5.**
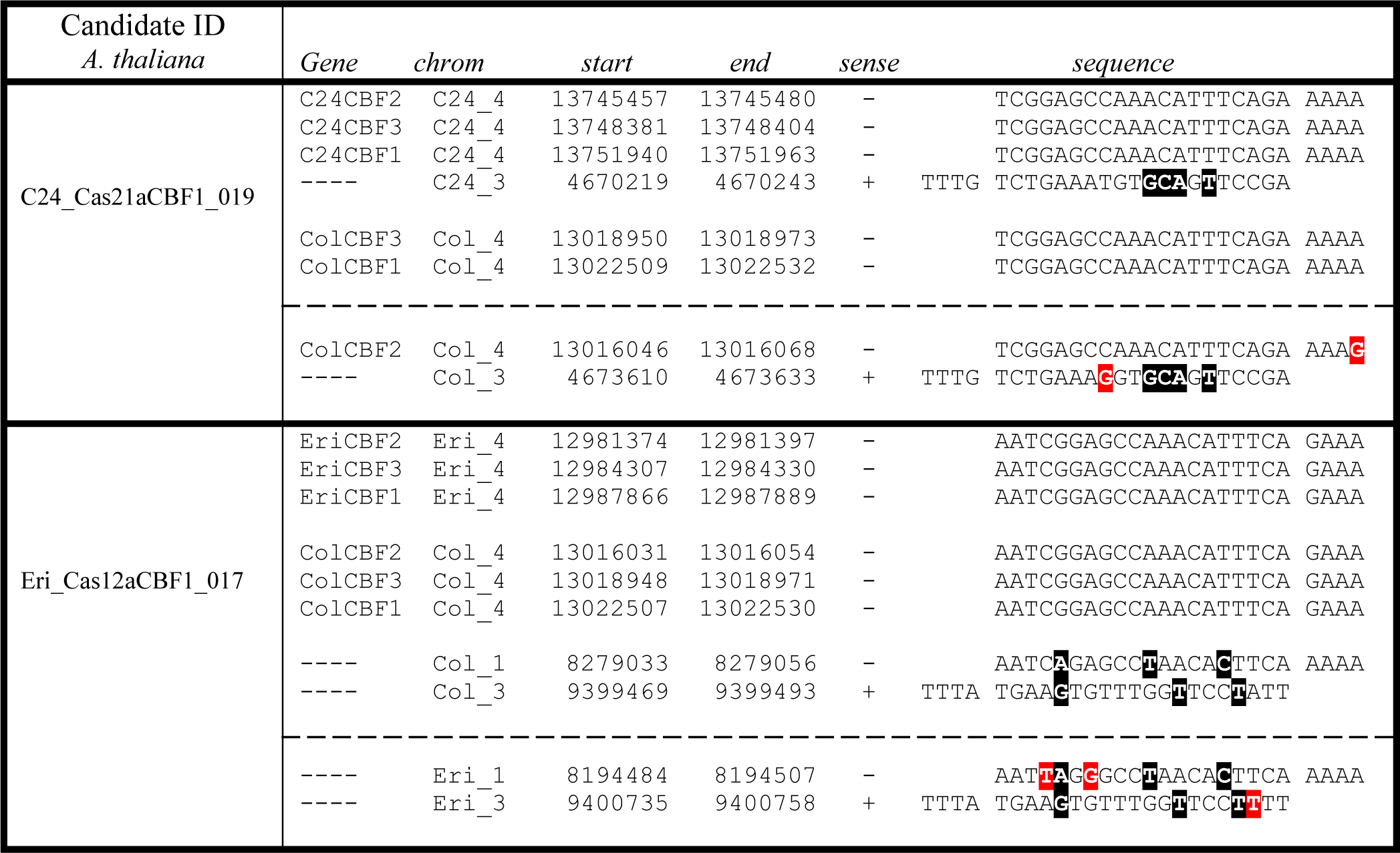
Intraspecies variability effect in the identification of targets and possible offtargets. For each example, upper file shows the targets and possible offtargets listed by ARES-GT (with thresholds L0 = 4 and L1 = 3) for each reference genome. SNPs differences between genomes that explain why some targets or offtargets are not detected are shown in lower file (separated by discontinuous line) as red boxes. Black boxes mark mismatches with candidates sequence.

## Conclusion

In summary, I have shown how the architecture of the ARES-GT tool (i) allows the selection of candidate sgRNAs that match multiple input query sequences for simultaneous editing of several members of gene families; (ii) contemplates the use of unmapped contigs apart from complete genomes; and (iii) can be used for the design of ecotype-specific CRISPR mutants. ARES-GT is available at GitHub (https://github.com/eugomin/ARES-GT.git).

---

**Supplemental Figure 1.**
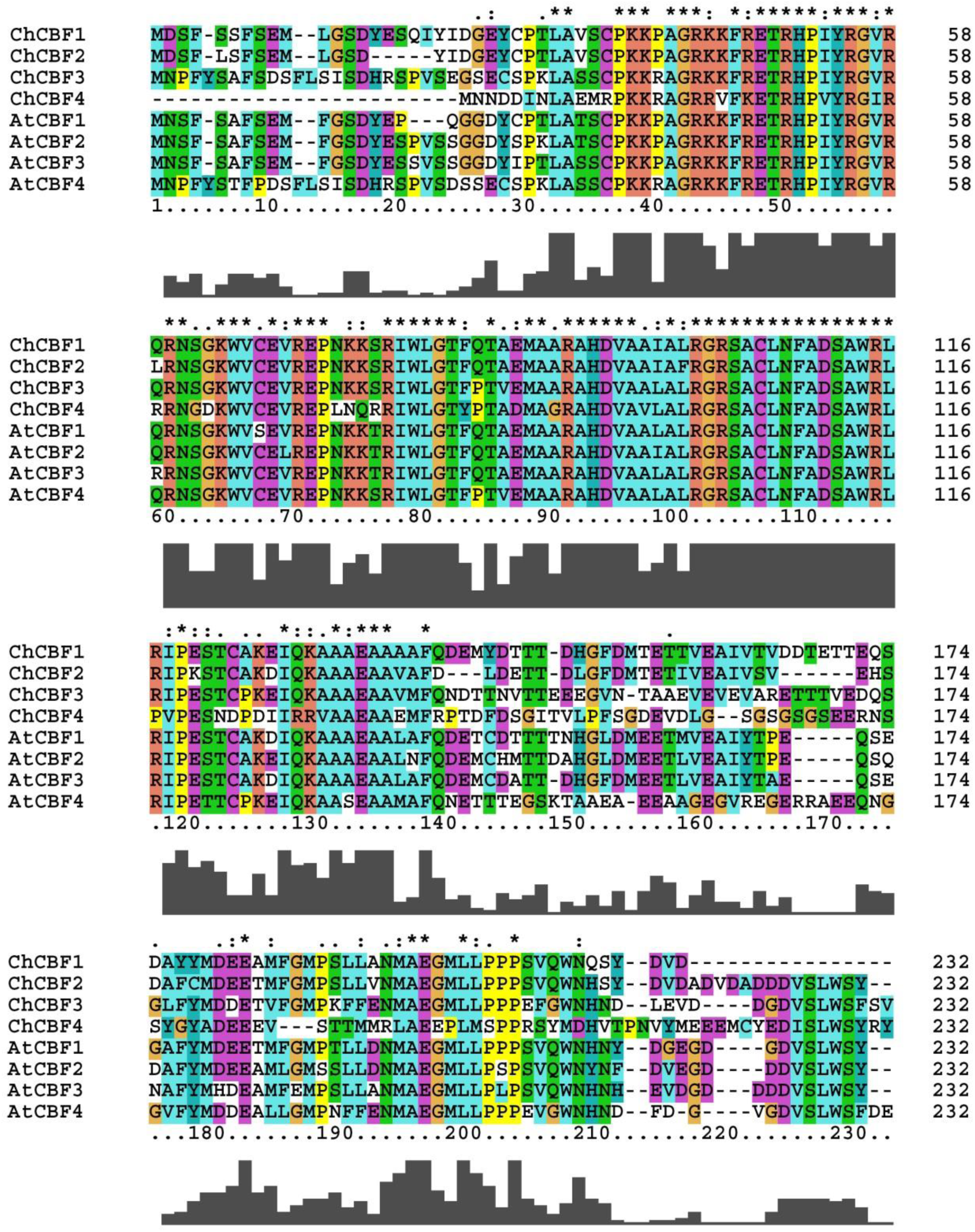
Alignment of CBF protein sequences from *Arabidopsis thaliana* and *Cardamine hirsuta*. Blast tool from *Cardamine hirsuta* genetic and genomic resource page (http://chi.mpipz.mpg.de/index.html) was used to find *AtCBF* homologs. Clustal Omega (Sievers et al., 2011) from EMBL-EBI tools (Madeira et al., 2019) was used for protein alignment.

